# A newly-identified IncY plasmid from multi-drug resistant *Escherichia coli* isolated from dairy cattle feces in Poland

**DOI:** 10.1101/2024.04.05.588223

**Authors:** Magdalena Zalewska, Aleksandra Błażejewska, Jan Gawor, Dorota Adamska, Krzysztof Goryca, Michał Szeląg, Patryk Kalinowski, Magdalena Popowska

## Abstract

Comprehensive whole-genome sequencing was performed on two multi-drug resistant *Escherichia coli* strains isolated from cattle manure from a typical dairy farm in Poland in 2020. The identified strains are resistant to beta-lactams, aminoglycosides, tetracyclines, trimethoprim/sulfamethoxazole, and fluoroquinolones. The complete sequences of the harbored plasmids revealed antibiotic-resistance genes (ARGs) located within many mobile genetic elements (e.g., insertional sequences or transposons), and genes facilitating conjugal transfer or promoting horizontal gene transfer. These plasmids are hitherto undescribed. Similar plasmids have been identified, but not in Poland. The identified plasmids carried resistance genes, including the tetracycline resistance gene *tet(A)*, aph family aminoglycoside resistance genes *aph(3”)-lb* and *aph(6)-ld*, β-lactam resistance genes *blaTEM-1, blaCTX-M-15*, sulfonamide resistance gene *sul2*, fluoroquinolone resistance gene *qnrS1*, and the trimethoprim resistance gene *dfrA14*. The characterized resistance plasmids were categorized into the IncY incompatibility group, indicating a high possibility for dissemination among the *Enterobacteriaceae*. While similar plasmids (99% identity) have been found in environmental and clinical samples, none have been identified in farm animals. This findings are significant within the One Health framework, as they underline the potential for antimicrobial-resistant *E. coli* from livestock and food sources to be transmitted to humans and vice versa. It highlights the need for careful monitoring and strategies to limit the spread of antibiotic resistance in the One Health approach.

## Introduction

*Escherichia coli* is a bacterial species found in the environment or colonizing the gastrointestinal tract of humans and animals. While most strains of *E. coli* are harmless commensals, some are pathogenic lineages capable of causing severe intestinal and extraintestinal diseases such as sepsis, urinary tract infections or meningitis (1). In addition, very strong evidence of gene exchange between strains found in food and clinics has been recorded (2). *E. coli* represents a major reservoir of antibiotic resistance genes (ARGs) that may be responsible for therapeutic failures in both human and veterinary medicine. Their numbers in *E. coli* isolates have increased during recent decades, with many of these ARGs being acquired by horizontal gene transfer (HGT). In the enterobacterial gene pool, *E. coli* may act as a donor and recipient of ARG, i.e. acquiring ARGs, such as the aminoglycoside-resistance gene *armA*, from bacteria and passing them to others.

HGT allows bacteria to adapt to unfavorable environmental conditions by acquiring sections of DNA; these genes can often include foreign DNA that can extend the ecological capabilities of the recipient. They can modify how the microorganism interacts with its host, give it an advantage over other microbes in the same environment, and, significantly, make it resistant to primary antibiotics by carrying ARGs on mobile genetic elements (MGEs). The roles of these MGEs are varied and in addition to conferring antibiotic resistance, they can facilitate digestion of various carbohydrate types, metal resistance, virulence, and catabolic processes beneficial for xenobiotic biodegradation in the environment (3). HGT is widespread and enables organisms from different taxonomic groups to share genetic material, effectively eroding the genetic distinctions between distinct phylogenetic groups and accounting for the substantial gene content variability among closely-related prokaryotes (4).

A key role in HGT is played by plasmids. These allow genetic material to be transferred at high rates through several pathways, primarily conjugation (including plasmid mobilization and conduction), but also through transduction, transformation, and vesiduction. Plasmids exhibit remarkable diversity with regard to size, copy number, G+C content, replication mechanisms, modes of transmission, DNA structure (either circular or linear), genetic content, and host spectrum. They harbor a wide array of traits that bolster their persistence and broaden the environmental niches that their hosts can occupy. Plasmids contain essential genes for both vertical and horizontal transmission, including those for replication and conjugation, and may also encode specialized systems to enhance transmission and stability. Many plasmids possess ’accessory’ genes that enrich the host’s ecological niche by enabling the degradation of toxic substances or introducing novel metabolic pathways. Notably, plasmids with virulence genes and ARGs play pivotal roles in the unchecked proliferation of bacterial pathogens within the One Health continuum (5, 6).

For instance, in 2003, the *armA* gene was identified on a conjugative plasmid in *Klebsiella pneumoniae* (7). Since then, this gene has been found in several enterobacteria, including *Proteus mirabilis, Serratia marcescens, Acinetobacter baumannii*, and *Pseudomonas aeruginosa* (8–10). In 2005, *armA* was found in *E. coli* isolated from farm animals, and its prevalence is still increasing. The dissemination of *armA* is facilitated by its location on the composite transposon Tn1548, which also harbors ARGs against sulfonamides located on self-transmissible plasmids belonging to several incompatibility groups. In 2017, a porcine *E. coli* isolate bearing *armA* was detected in Italy; it was multidrug-resistant (MDR), harboring the *blaCMY-2*, *blaOXA-181*, and *mcr-1* genes additionally (11). This illustrates how introducing a new gene coding for resistance, through adapting to the environment, should be a matter for concern. Genes pose an even greater threat than antibiotics because when they emerge, they cannot be removed from the environment (12).

ESBL (extended-spectrum β-lactamases)/AmpC genes have been found to be highly prevalent in commensal *E. coli* isolated from fecal samples of various food-producing animals, and many of them have been found to be located on plasmids. The TEM- and SHV-ESBLs phenotypes were predominant until 2000, then CTX-M-ESBLs emerged. After the 2000s, this phenotype has been mostly identified in commensal and pathogenic ESBL-producing *E. coli* isolates of humans and animals. Carbapenemases have been rarely identified in *E. coli* strains of animal origin as they are very rarely prescribed in veterinary medicine and hence exert very weak selective pressure; however, the prevalence of carbapenemase-producing *E. coli* in animals worldwide has recently increased (VIM, NDM, IMP, OXA, and KPC phenotypes). Even though the occurrence of carbapenemase-producing *Enterobacteriaceae* identification in animals is marginal, it still poses a significant threat to human medicine (13, 14).

Several plasmid-encoded resistance mechanisms to quinolones have been identified in animal- and human-related sources, such as Qnr-like proteins, which protect DNA from binding to quinolone, the acetyltransferase that modifies certain fluoroquinolones, and active efflux pumps. These resistance determinants do not confer high resistance to antimicrobial quinolones, or fluoroquinolones, but rather confer reduced susceptibility to them (15). MGE often facilitates the rapid spread of resistance to different hosts: indeed, insertion sequences (ISs), transposons, integrons and conjugative plasmids can all move DNA between replicons or cells (16).

As a consequence of the high selective pressure imposed by the widespread use of tetracyclines, many bacteria, including *E. coli*, have developed or acquired MGE resistance to tetracycline: nine tetracycline efflux genes *tetA, tetB, tetC, tetD, tetE, tetG, tetJ, tetL*, and *tetY*, two tetracycline resistance genes encoding ribosomal protective proteins *tetM* and *tetW*, and one gene coding for an oxidoreductase that inactivates tetracyclines *tetX* have been identified in *E. coli*. The major mechanisms of tetracycline resistance encountered in *E. coli* of animal origin cover (i) the active efflux by proteins of the major facilitator superfamily and (ii) ribosome protection (17).

Also, aminoglycoside resistance genes, which can be classified as target modification (*armA, rmtA, nmpA*) or antimicrobial inactivating (most prevalent *strA, strB*) enzymes, have been found in *E. coli* mobilized on resistance plasmids. These provide resistance against only the most extensively-studied antimicrobial groups applied for both human and animal medicines; more mechanisms have been reviewed in detail by Poirel et al. (17).

Generally, ARGs in *E. coli* are considered a major global challenge in human and animal medicine and must be considered a real public health concern. *E. coli* infections in animals are treated with a range of antimicrobials: ampicillin, streptomycin, sulfonamides, or oxytetracyclines are commonly used to treat bovine mastitis and newborn diarrhea, with broad-spectrum cephalosporins and fluoroquinolones also possibly being used. Animal feces are important reservoirs of human pathogens such as pathogenic *E. coli*, *Salmonella,* or *Listeria* (18). The co-occurrence of various ARGs originating in animal intestines and microbial human pathogens, together with MGEs, creates perfect conditions for disseminating antibiotic resistance through the acquisition of ARGs by bacteria (19).

The present study examines *E. coli* strains extracted from the feces of dairy cattle, specifically from intensive breeding facilities. The antimicrobial susceptibility and plasmid profiles of the strains were tested, and the most common strains selected for sequencing using the NovaSeq 6000 (Illumina) and MinIon (Oxford Nanopore) platforms. As a result, the study presents the full structure of the plasmids and the genetic information contained therein.

## 2. Materials and Methods

### 2.1 Isolation of *E. coli* strains

Samples of dairy cattle manure were collected from a Polish dairy farm employing an intensive rearing system. The animals were highly-productive Polish Holstein-Friesian dairy cows of the Black and White variety. The herd was located in the central part of Poland in the Masovian district. The manure sampling procedure is described in more detail by Zalewska et al. (20). To isolate antibiotic-resistant *E. coli* strains, the waste sample was enriched in Luria Bertani broth by adding 1 g of feces to 9 mL of liquid medium. The cultures were incubated at 30 °C and 37 °C for 24 hours and 48 hours. After incubation, the bacterial suspensions were diluted (10^−1^, 10^−2^, and 10^−3^), and 100 µL of undiluted sample and each dilution were plated onto Eosin Methylene Blue Agar and MacConkey Agar: one set was plated onto the two media supplemented with imipenem (16 mg/L) to confirm a carbapenem resistance phenotype, and another set on the media supplemented with cefotaxime (4 mg/L) to test for an Extended-spectrum beta-lactamase phenotype (21). Each isolation was performed as three biological replicates, followed by three technical ones. All selective media were purchased from Biomaxima (Poland). Antibiotics were purchased from Sigma-Aldrich (MERCK, USA). The plates were incubated at 30 °C and 37 °C for 24 hours to select for animal pathogens and environmental strains. Next, approximately 24 to 48 colonies showing the morphology of the intended bacteria were taken and applied to three consecutive streaks to get pure colonies. The pure cultures were stored in PBS/glycerol stocks (20% v/v) for further analysis.

#### The susceptibility profiles of *E. coli* strains

The antibiotic susceptibility profiles of the isolated strains were determined by The Kirby– Bauer test (22). The phenotypic characterization was based on the following antimicrobial discs: imipenem (IMP 10 µg), ciprofloxacin (CIP 5 µg), cefotaxime (CTX 5 µg), and gentamicin (CN 10 µg) (OXOID, USA). The inhibition zones were measured according to EUCAST recommendations, i.e. after incubation for 18 hours at 37 °C.

Strains with different antimicrobial susceptibility profiles, characterized by a difference in bacterial growth inhibition zone diameter of approximately +/− 2 mm around at least one antibiotic disk, were considered non-repetitive and chosen for further research. The isolated strains were confirmed as *E. coli* by MALDI-TOF MS/MS (matrix-assisted laser desorption/ionization system equipped with a time-of-flight mass spectrometer) (23). The analyses were performed in an external medical laboratory, ALAB Laboratoria Sp. z o. o. (Poland), according to a standard diagnostic procedure. Identification was performed by aligning the peaks to the best matching reference data, with the resulting log score classified as follows: ≥2.3 indicates a highly probable species, 2.0 – 2.3 a certain genus and probable species, 1.7 – 2.0 a probable genus, and <1.7, non-reliable identification. Antibiotic susceptibility testing was then performed with Vitek2 Compact equipment (BioMerieux, France) (24), with strains being classified as sensitive (S), resistant (R), or intermediate (I) based on the results. The AST-N331 susceptibility card was used. Strains with the same resistance profile were considered clones.

The plasmid profile of the non-repetitive strains was determined in three ways: 1) isolation of plasmid DNA with a commercially-available kit – Plasmid Mini (A&A Biotechnology, Poland); 2) rapid alkaline lysis for the isolation of plasmid DNA (25); 3) plasmid visualization by Eckhardt electrophoresis (26). Examples of plasmid DNA with similar profiles were additionally differentiated by digestion with restriction enzymes BamHI and HindIII (NEB, USA). The digestion was performed according to the manufacturer’s recommendation. Depending on the product size, the isolation and digestion effect was visualized with electrophoresis in 1.5% to 2% agarose. Strains with different profiles were chosen for sequencing.

### 2.3 DNA extraction and sequencing

Genomic DNA was isolated from bacterial strains using Genomic Mini (A&A Biotechnology, Gdynia, Poland). The DNA concentration was measured using a Qubit fluorometer and a dsDNA High Sensitivity Assay Kit (Thermo Fisher Scientific, USA), and purity was determined by measuring the A260/A280 absorbance ratio with a NanoDrop spectrophotometer (Thermo Fisher, USA). Only samples with concentrations higher than 10 ng/µL and an A260/A280 ratio ranging from 1.8 to 2.0 were analyzed. DNA samples were stored at −20 °C for further use. The DNA samples were isolated in triplicate.

### 2.4 Genomic DNA sequencing on Oxford Nanopore MinION

Samples composed of 500 ng of isolated DNA were taken for library preparation. They were first mechanically fragmented using a syringe-based method (0.4 x 20 mm needle, 1 ml glass syringe) in a volume of 200 µl. The fragmented samples were purified and concentrated to a volume of 13 µl using the SPRI method (Solid Phase Reversible Immobilization) with Kapa Pure Beads (Roche, cat. no. 07983298001). Elution was performed for 10 minutes at 37°C. Libraries were constructed using the manufacturer’s protocol of Native Barcoding Kit 24 with genomic DNA (SQK-NBD112.24, Version: NBE_9134_v112_revE_01Dec2021) using a NEB Blunt/TA Ligase Master Mix (New England Biolabs, cat. no. M0367), a NEBNext Quick Ligation Reaction Buffer (New England Biolabs, cat. no. B6058) and a NEBNext Companion Module for Oxford Nanopore Technologies (New England Biolabs, cat. no. E7180S). Sequencing was performed on the NanoPore MINIon MkB1 instrument using the R10.4 version of reagents (cat. no. FLO-MIN112) following the manufacturer’s protocol. Basecalling was performed with guppy (version 6.1.2) with --flowcell FLO-MIN112 --kit SQK-NBD112-24 options.

### 2.5 Genomic DNA sequencing on Illumina NovaSeq 6000

The DNA samples were fragmented using the S220 Focused-ultrasonicator (Covaris) to an mean size of approximately 300 bp (Duty Factor: 10%, Peak Incident Power: 140, Cycles per Burst: 200, Treatment Time: 80s). Subsequently, libraries were constructed using the KAPA Hyper Prep Kit (Roche, cat. no. 07962363001) and TruSeq DNA UD Indexes adapters (Illumina, cat. no. 20020590) according to the manufacturer’s recommendations, starting with 100 ng of material. Five cycles of PCR amplification were performed for enrichment. Fragment size selection was carried out using Kapa HyperPure Beads (Roche, cat. no. 07983298001) in a two-step purification with a sample-to-reagent volume ratio of 1) step - 1:0.75, 2) step - 1:0.85. Sequencing was performed on an Illumina NovaSeq 6000 instrument using the NovaSeq 6000 S1 Reagent Kit v1.5 (200 cycles) (Illumina, cat. no. 20028317), in a pair-end 2x100 cycle mode, using the standard procedure recommended by the manufacturer with a 0.5% addition of the PhiX control library (Illumina, cat. no. FC-110-3001).

### 2.6 Genome assembly and data analysis

Sequence quality metrics for Illumina data were assessed using FASTQC v.0.12.0 (http://www.bioinformatics.babraham.ac.uk/projects/fastqc/) (27) and short reads were quality trimmed using fastp v.0.23.2 (28). Nanopore reads were quality filtered using NanoFilt v.2.8.0 (reads shorter than 1kb and QV<12 were discarded) (29). After residual adapter removal using Porechop v.0.2.4 (https://github.com/rrwick/Porechop) the dataset was quality checked using NanoPlot v. 1.41.6 (29).

Long-read assembly was performed using Trycycler v.0.5.3 pipeline (30). In brief, nanopore reads were initially assembled using four long read assemblers: flye v2.9, unicycler v0.4.8, Raven v1.8.1 and miniasm v0.3-r179. The long-read assemblies were then reconciled and circularized, and final consensus was generated followed by polishing with medaka (https://github.com/nanoporetech/medaka). Long-read assembled contigs were further polished using polypolish (https://github.com/rrwick/Polypolish) (30) and POLCA (31). All of the possible sequence errors and misassemblies were further manually corrected using SeqMan software (DNAStar) to obtain a complete nucleotide sequence of bacterial genome.

The assembled genomic sequences were annotated using Bakta (32). Plasmid replicon sequences were searched against the PLSDB database ((https://ccb-microbe.cs.uni-saarland.de/plsdb/). ARGs were detected and localized in the genomic sequences using abricate (https://github.com/tseemann/abricate). MGEs were identified using mobileelementfinder (33) and the ARG-associated mobilome was characterized using VRprofile2 (34).

The plasmid sequences were visualized using the SnapGene Viewer software (SnapGene® software from Dotmatics; available at snapgene.com). The linear plasmid sequences were compared using Easyfig software (35).

## 3. Results

### *E. coli* strain isolation

In total, 27 bacterial strains demonstrating growth indicative of *E. coli* were found using selective media; more specifically, 16 strains were isolated using EMB Agar and 11 via MacConkey Agar (Supplementary Table 1). Among the isolated bacteria, 12 were identified as *E. coli* through MALDI-TOF MS/MS analysis, of which four strains were isolated on EMB agar and eight on MacConkey agar. All bacterial strains were isolated on agar plates enriched with cefotaxime; however, no strains exhibited resistance to imipenem.

The antimicrobial susceptibility profiles of all strains were comprehensively determined utilizing the Vitek 2 Compact system (Supplementary Table 2). Isolates that demonstrated analogous antimicrobial susceptibility profiles were aggregated, and their plasmid profiles were elucidated. Following the comparative analysis of plasmid profiles within these aggregates, two bacterial strains exhibiting potentially unique plasmid configurations were selected for total plasmid DNA sequencing.

### Genome sequencing and assembly

The genomes of the investigated strains were sequenced using a hybrid approach. Oxford Nanopore long-read technology (MINIon platform) was used to obtain as complete scaffolds as possible. The quality of the obtained sequences was then improved with Illumina short-read technology (2x100 nt in pair-end mode, NovaSeq 6000, approximate 100x coverage). The bacterial chromosome was assembled for both samples, resulting in an approximately 5Mb contig. Four additional significant contigs were assembled for “kr1” and three for “kr2 (Table 1), and these analyzed in detail.

**Table 1.** Plasmids identified in studied *E. coli* strains. Those harboring ARGs are marked in gray.

| Strain no. | Origin | Plasmid no. | Plasmid name in our collection | Size [bp] |
| --- | --- | --- | --- | --- |
| kr1 | cattle manure | 1 | pECmdr1.1 | 103976 |
|  |  | 2 | pECmdr1.2 | 6647 |
|  |  | 3 | pECmdr1.3 | 6466 |
|  |  | 4 | pECmdr1.4 | 3374 |
| kr2 | cattle manure | 1 | pECmdr2.1 | 103976 |
|  |  | 2 | pECmdr2.2 | 6647 |
|  |  | 3 | pECmdr2.3 | 6466 |

### Plasmids

Seven plasmids belonging to two bacterial isolates were identified (Table 1). The complete genomes of *E. coli* are deposited as a part of ‘INART Escherichia coli genomes’ bioproject (Accession no. PRJNA942482). Two plasmids have more than 100,000 bp and may be classified as megaplasmids. These two identified plasmids harbored different ARGs and MGEs (Table 2). Plasmids that do not confer antibiotic resistance will be characterized and deposited further, as a part of an additional project.

**Table 2.**
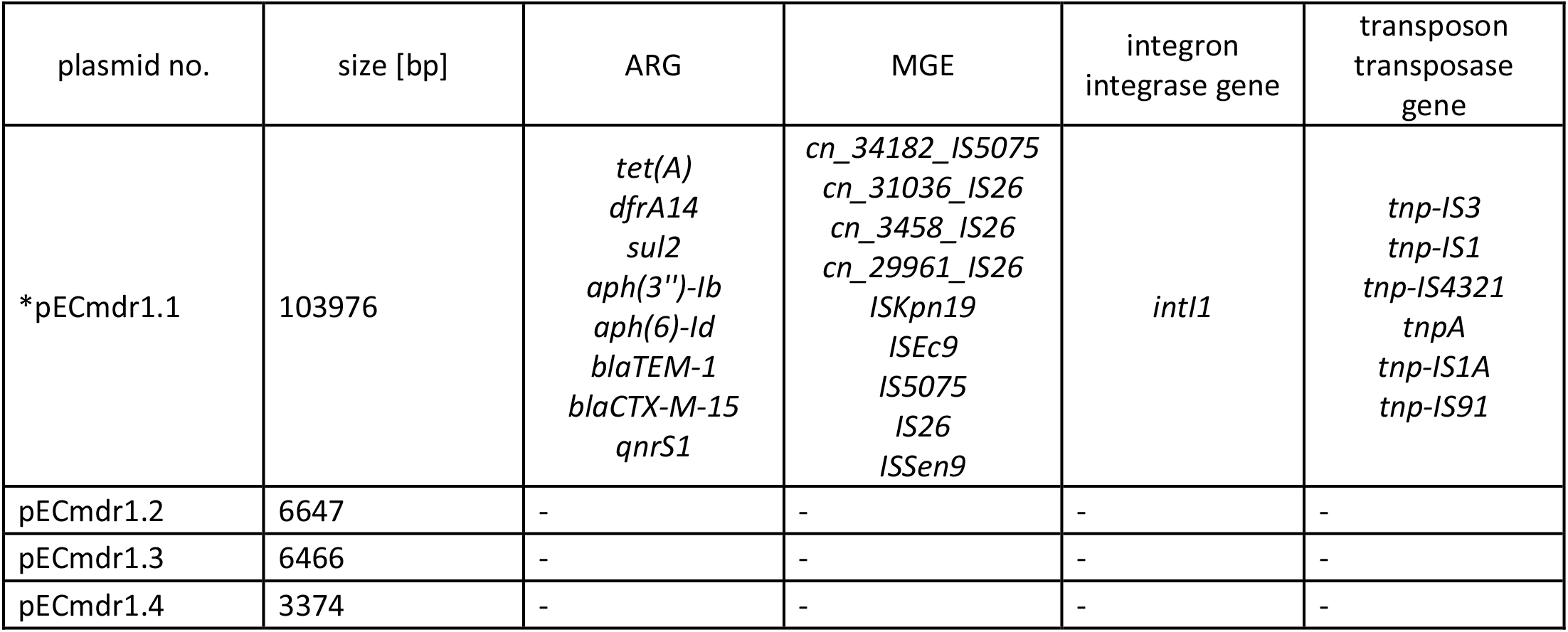
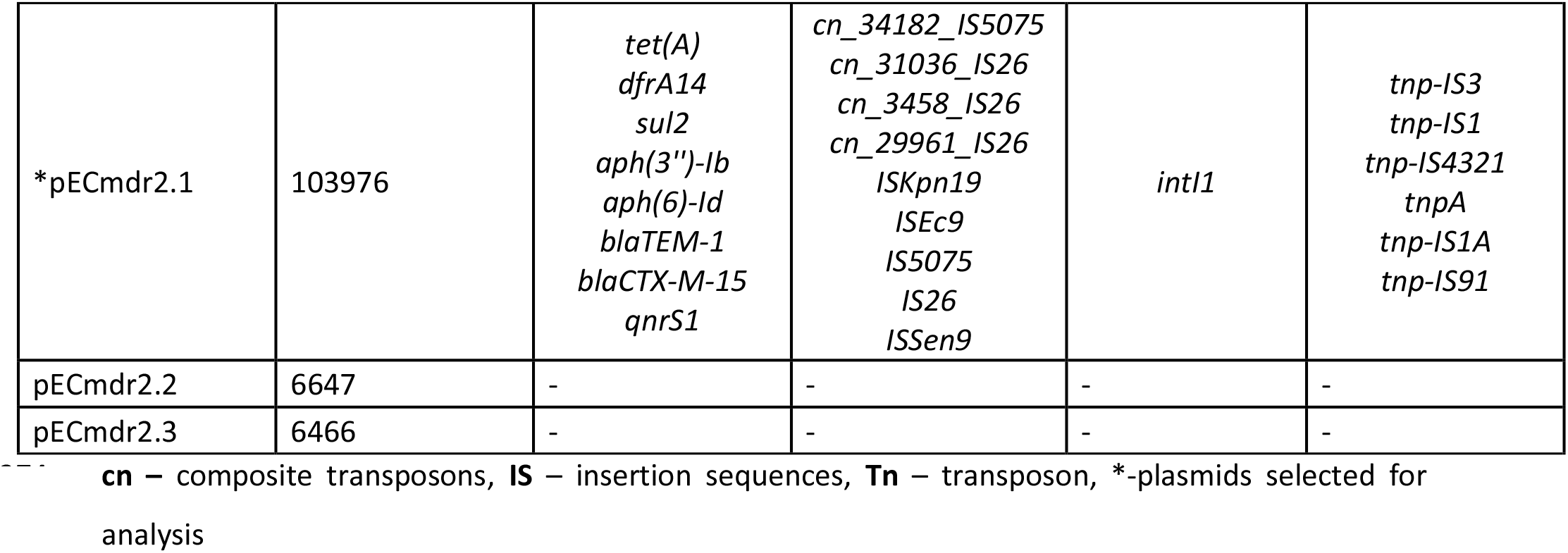
ARGs, MGEs, integron integrase genes, transposon transposase genes, conjugal transfer system genes and heavy metal resistance genes.

| plasmid no. | size [bp] | ARG | MGE | integron integrase gene | transposon transposase gene |
| --- | --- | --- | --- | --- | --- |
| *pECmdr1.1 | 103976 | *tet(A)*<br>*dfrA14*<br>*sul2*<br>*aph(3'')-Ib*<br>*aph(6)-Id*<br>*blaTEM-1*<br>*blaCTX-M-15*<br>*qnrS1* | *cn_34182_IS5075*<br>*cn_31036_IS26*<br>*cn_3458_IS26*<br>*cn_29961_IS26*<br>*ISKpn19*<br>*ISEc9*<br>*IS5075*<br>*IS26*<br>*ISSen9* | *intI1* | *tnp-IS3*<br>*tnp-IS1*<br>*tnp-IS4321*<br>*tnpA*<br>*tnp-IS1A*<br>*tnp-IS91* |
| pECmdr1.2 | 6647 | - | - | - | - |
| pECmdr1.3 | 6466 | - | - | - | - |
| pECmdr1.4 | 3374 | - | - | - | - |
| *pECmdr2.1 | 103976 | *tet(A)*<br>*dfrA14*<br>*sul2*<br>*aph(3'')-Ib*<br>*aph(6)-Id*<br>*blaTEM-1*<br>*blaCTX-M-15*<br>*qnrS1* | *cn_34182_IS5075*<br>*cn_31036_IS26*<br>*cn_3458_IS26*<br>*cn_29961_IS26*<br>*ISKpn19*<br>*ISEc9*<br>*IS5075*<br>*IS26*<br>*ISSen9* | *intI1* | *tnp-IS3*<br>*tnp-IS1*<br>*tnp-IS4321*<br>*tnpA*<br>*tnp-IS1A*<br>*tnp-IS91* |
| pECmdr2.2 | 6647 | - | - | - | - |
| pECmdr2.3 | 6466 | - | - | - | - |

**cn** – composite transposons, **IS** – insertion sequences, **Tn** – transposon, \*-plasmids selected for analysis

Fig. 1 and 2 present the linear and circular plasmid structures of the bacteria, with particular regard to ARGs, MGEs, conjugal transfer genes (TRA), transposon transposase genes (TNF), heavy metal resistance genes, replication system (REP), integron integrase, partitioning system elements (PAR).

**Figure 1.**
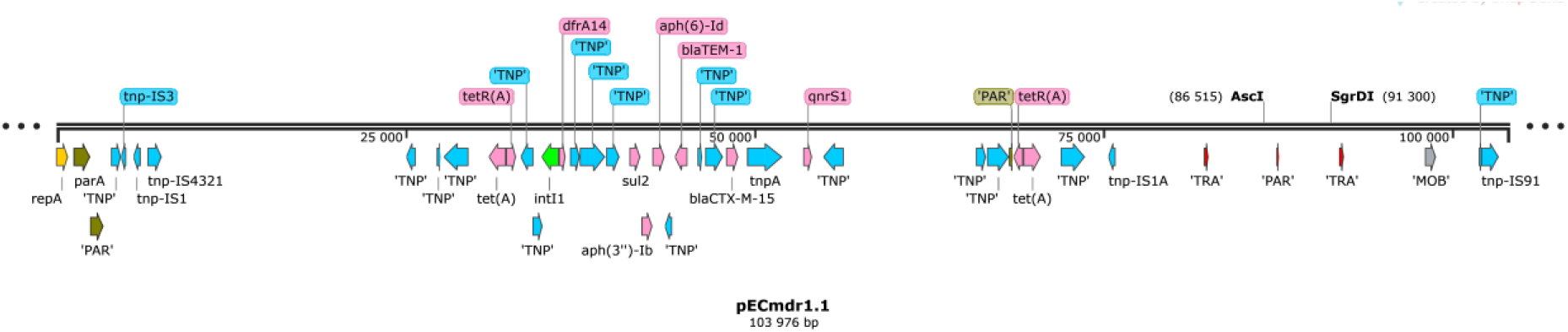
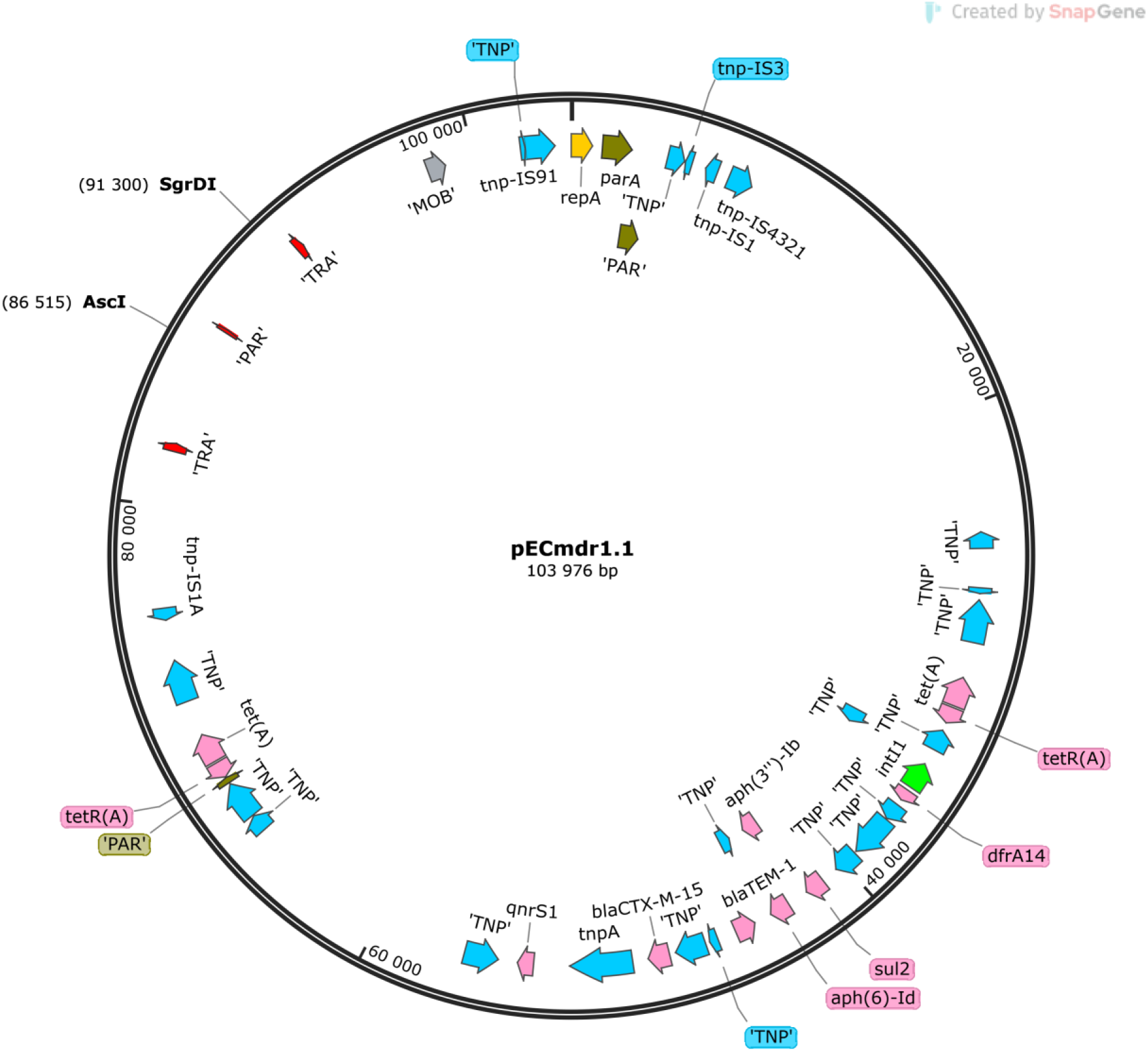
Linear and circular representation of pECmdr1.1 plasmid (pink-ARG, red-conjugal transfer genes TRA, blue-transposon transposase genes TNP, brown-heavy metal resistance genes, yellow- replication system REP, green-integron integrase, khaki-partitioning system PAR, gray-mob system MOB, plum-pilus system PIL; quotation marks indicate that the function of the gene has been assigned based on the homology of the protein it encodes)

**Figure 2.**
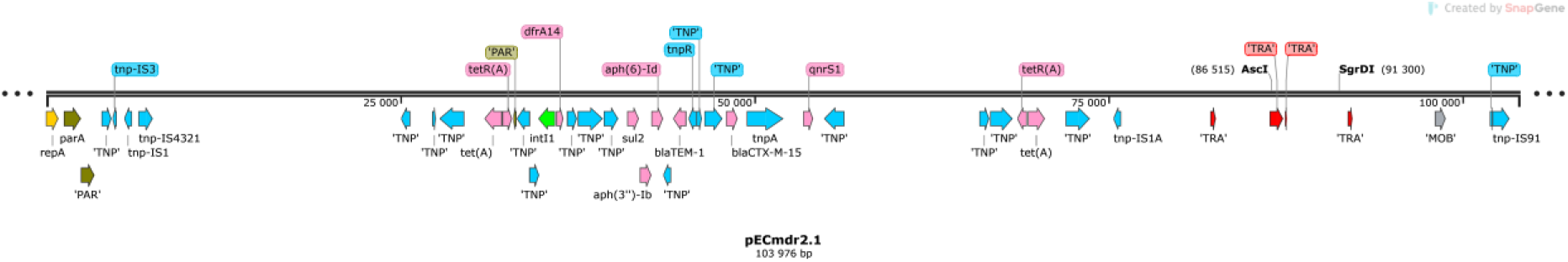
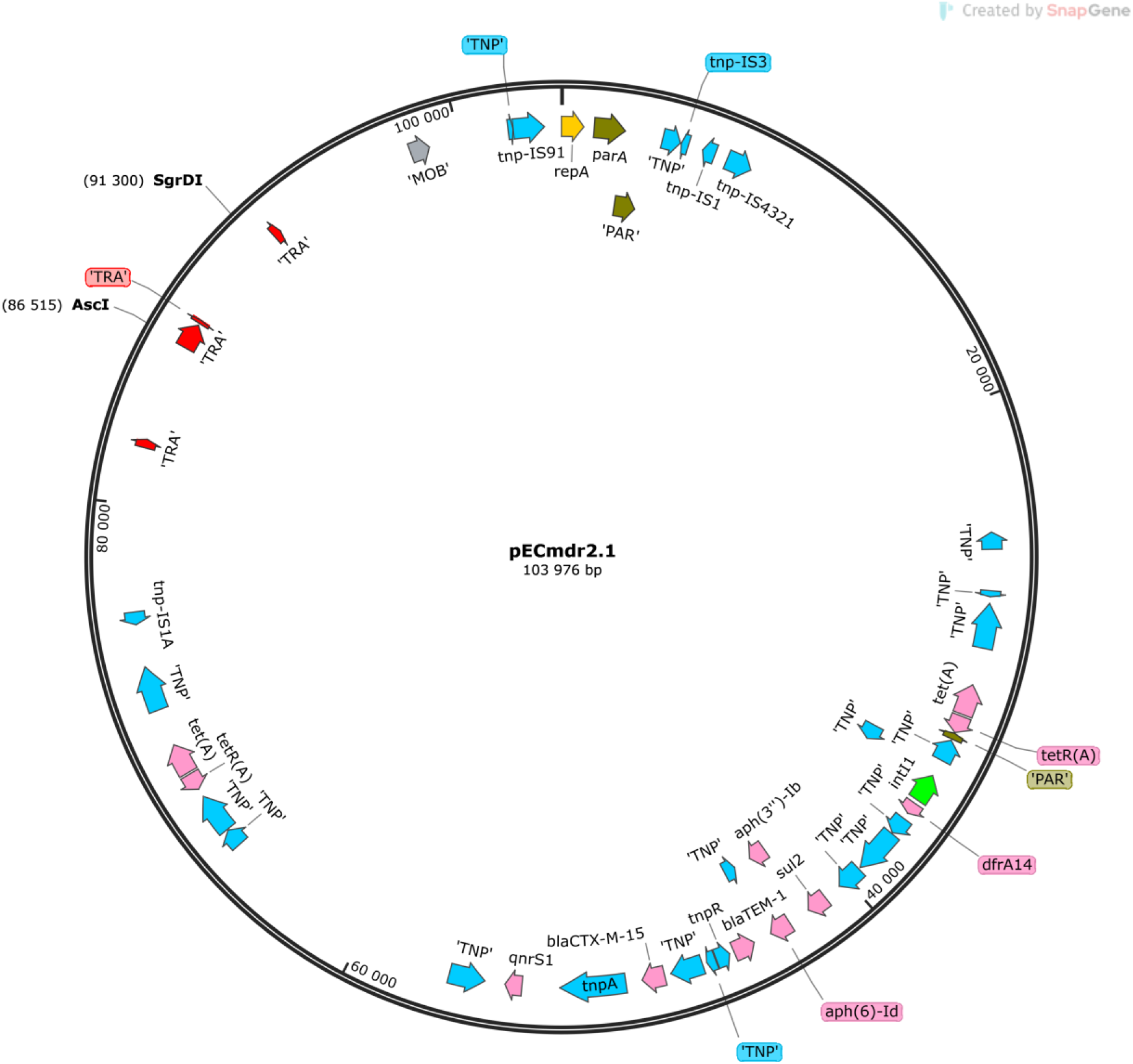
Linear representation of pECmdr2.1 plasmid (pink-ARG, red-conjugal transfer genes TRA, blue-transposon transposase genes TNP, brown-heavy metal resistance genes, yellow-replication system REP, green-integron integrase, khaki-partitioning system PAR, gray-mob system MOB, plum- pilus system PIL; quotation marks indicate that the function of the gene has been assigned based on the homology of the protein it encodes

Although these plasmids are similar in size and carry genes, particularly ARG, MGE, PAR system, TNP system, TRA system, or REP system, they differ structurally (Fig. 3).

**Figure 3.**
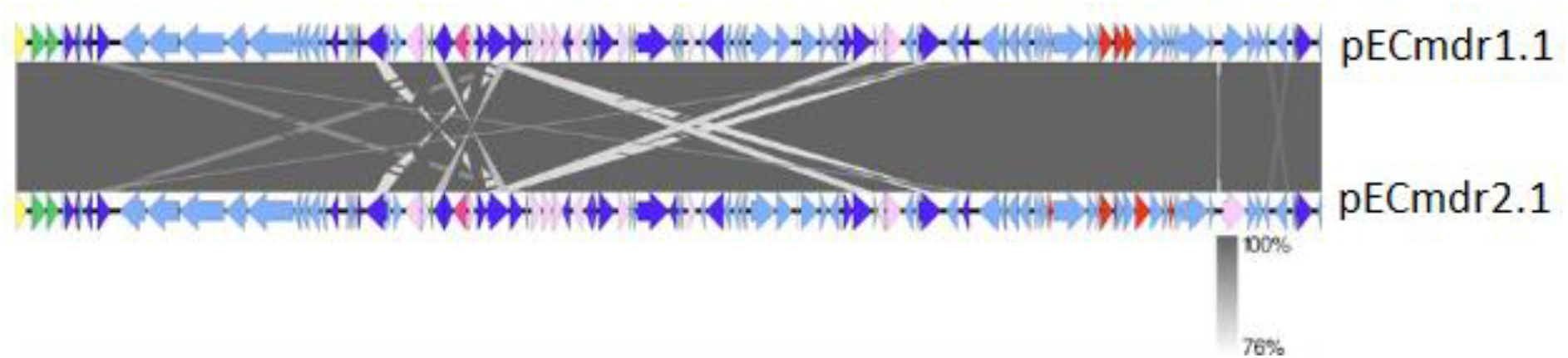
Comparison between pECmdr1.1 and pECmdr2.1 (linear map showing the genetic structure of circular plasmids) (yellow-replication system, red-conjugal transfer system, green-partitioning system, violet-ARG, blue-transposase genes, pink-integron integrase, pale blue-other genes)

A detailed analysis of bacterial plasmids sequences deposited in the PLSDB plasmid database (https://ccb-microbe.cs.uni-saarland.de/plsdb/) (36, 37) found the plasmids identified during the study to be closely similar with previously-deposited ones, albeit with some differences. The pECmdr1.1 plasmid has 29 similar hits, while pECmdr2.1 has 29 (identity threshold 0.99).Sequence analysis indicates that both plasmids belong to the incY incompatibility group. Neither plasmid has previously been reported in Poland, but they have been identified in many other countries. Plasmids similar to pECmdr1.1 and pECmdr2.1 have been found only in the water environment and human clinical samples: not in cattle (Table 3).

**Table 3.**
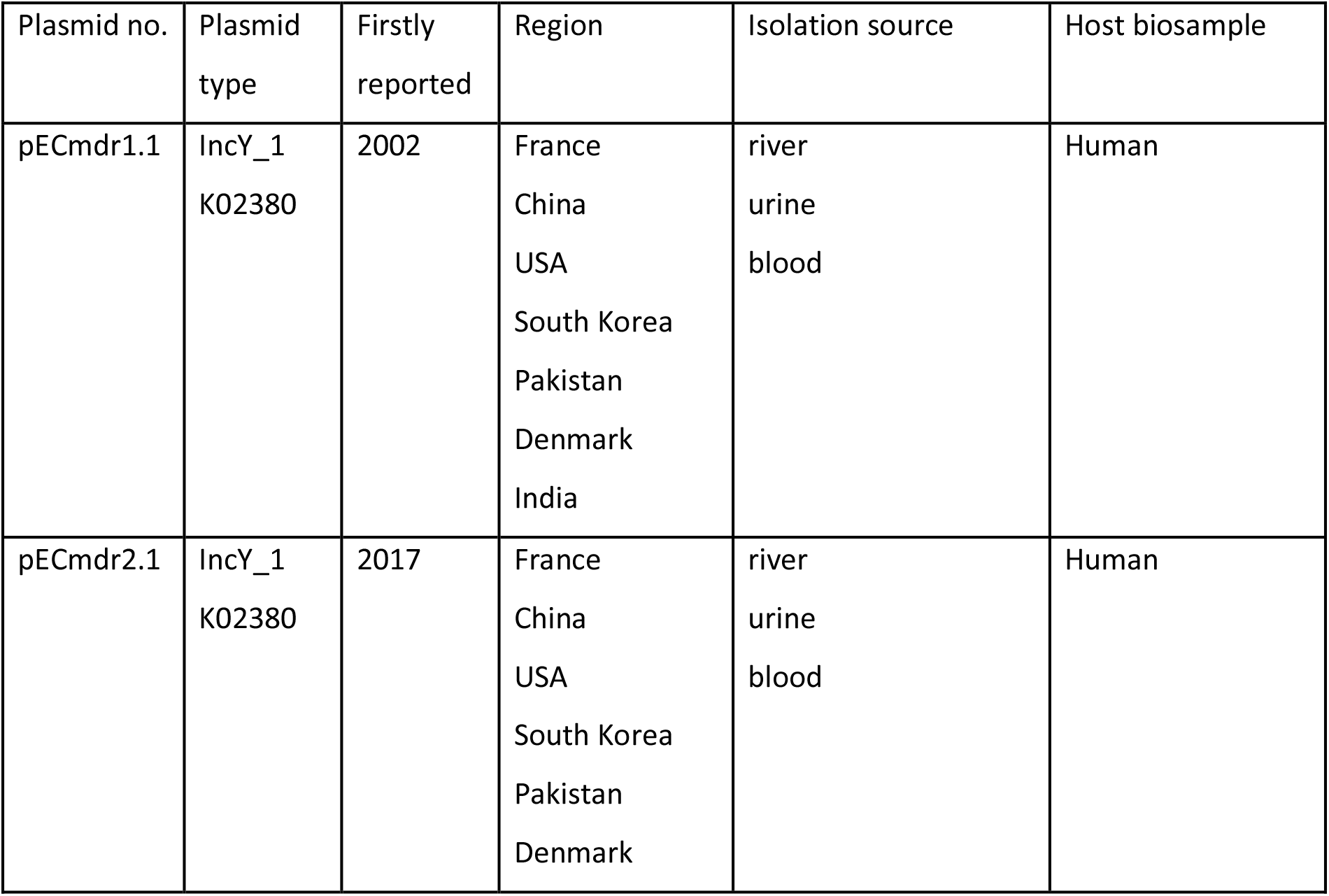
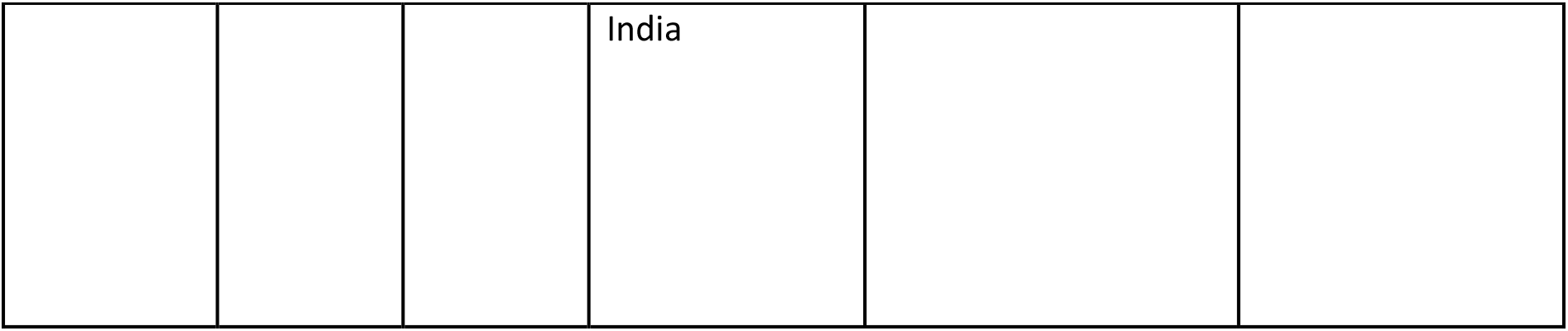
Region, isolation source, and host biosample of similar plasmids (plasmid identity 0.99).

**Table 4.** The host range for similar plasmids (plasmid identity 0.99)

| Plasmid similar to | identified in the following bacteria: |
| --- | --- |
| pECmdr1.1 | *Escherichia coli*<br>*Klebsiella pneumoniae*<br>*Salmonella enterica subsp. enterica* |
| pECmdr2.1 | *Escherichia coli*<br>*Klebsiella pneumoniae*<br>*Salmonella enterica subsp. enterica* |

By aligning the ARG-coding sequences with the defined and described MGE, it was possible to identify the ARGs located within the MGE, which were situated on the identified plasmids (Fig. 4A-D).

**Figure 4.**
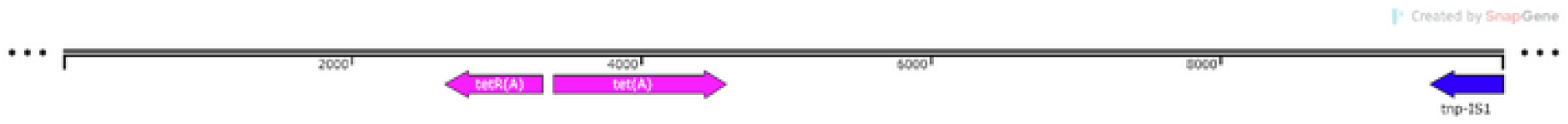
A. pECmdr1.1 ISclusterTn (9949 bp)

**Figure 4.**
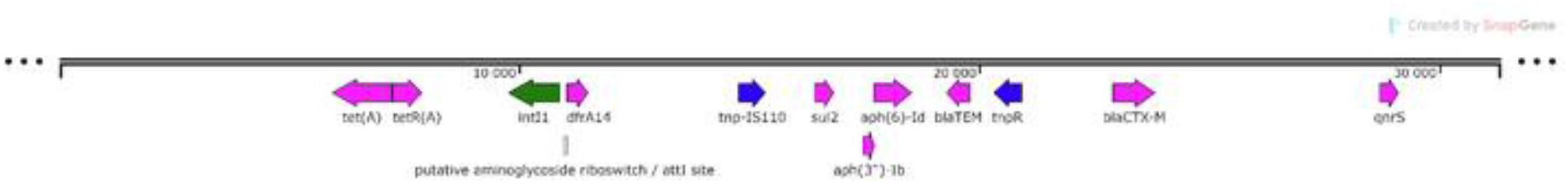
B. pECmdr1.1 ISclusterTn (31306 bp)

**Figure 4.**
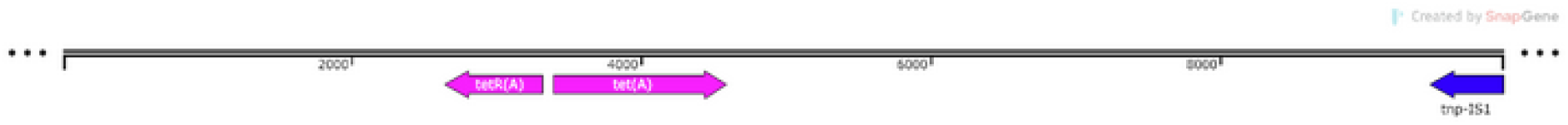
C. pECmdr2.1 ISclusterTn (9949 bp)

**Figure 4.**
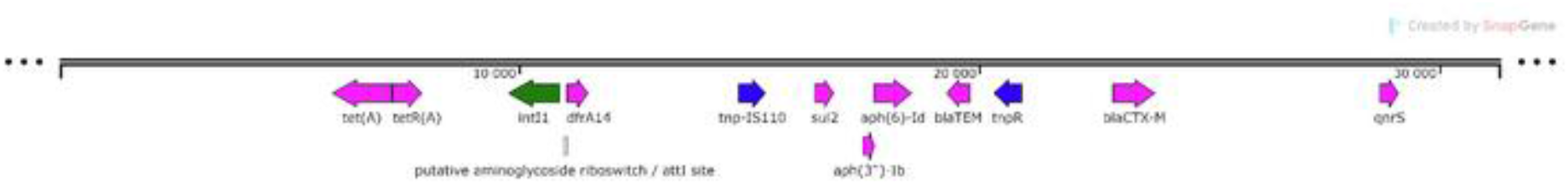
D. pECmdr2.1 ISclusterTn (31306 bp)

## Discussion

The spread of antimicrobial resistance is recognized as a key global challenge to human health (38). *E. coli* in particular is known to cause many infections, and to be frequently associated with drug resistance, especially against cephalosporins (ESBL phenotype) (17). The risk related to infection caused by carbapenem-resistant, ESBL-producing *Enterobacteriaceae* has been highlighted by the World Human Organization as critical (21). Current research in Microbiology suggests that the diversity of ARGs in plasmids can be linked to the varied environments where bacteria thrive. Studies have shown that certain ARGs, like *tet(A)*, are commonly found in both clinical and environmental isolates (CARD) (39), indicating a widespread distribution and a potential for cross-environment transfer.

The high diversity of bacterial ARGs and the varied health risks associated with bacterial human pathogens require proper environmental risk assessment. ARGs from anthropogenic pollutants have considerable potential to influence environmental resistomes and to be acquired by human pathogens; as such they constitute a significant environmental and health threat. Human-animal-shared mobile ARGs have been identified in human, chicken, pig and cattle guts, and studies suggest they may confer resistance to as many as six major antibiotic classes (tetracyclines, aminoglycosides, macrolide-lincosamide-streptogramin B, chloramphenicol, beta-lactams, and sulphonamides) at the same time. Additionally, there is a shared resistome between soil bacteria and human pathogens, and it is suggested that soil microbiota act as ARG reservoirs for human and animal pathogens (40, 41).

Our present findings identified two very similar plasmids harboring ARGs against at least six antimicrobial groups, making the identified bacterial strains multi-drug resistant (MDR). Both plasmids harbored the tetracycline resistance gene *tet(A)* (TetA is a tetracycline efflux pump found in many species of Gram-negative bacteria), the trimethoprim resistance gene *dfrA14* (dfrA14 is an integron-encoded dihydrofolate reductase found in Gram-negative bacteria), the sulfonamide resistance gene *sul2* (Sul2 is a sulfonamide resistant dihydropteroate synthase of Gram-negative bacteria, usually found on small plasmids), the aminoglycoside resistance gene *aph(3’’)-Ib* (APH(3’’)- Ib is an aminoglycoside phosphotransferase encoded by plasmids, transposons, integrative conjugative elements and chromosomes in *Enterobacteriaceae* and *Pseudomonas* spp.), the aminoglycoside resistance gene *aph*(*6*)*-Id* (APH(6)-Id is an aminoglycoside phosphotransferase encoded by plasmids, integrative conjugative elements and chromosomal genomic islands in *Enterobacteriaceae*, *Providencia alcalifaciens*, *Pseudomonas* spp., *Vibrio cholerae*, *Edwardsiella tarda*, *Pasteurella multocida* and *Aeromonas bestiarum*), *blaTEM-1* (beta-lactam resistance gene; TEM-1 is a broad-spectrum beta-lactamase found in many Gram-negative bacteria; confers resistance to penicillins and first generation cephalosphorins), the beta-lactam resistance gene *blaCTX-M-15* (CTX-M-15 is a beta-lactamase found in the *Enterobacteriaceae* family) and the quinolone-resistance gene *qnrS1* (QnrS1 is a plasmid-mediated quinolone resistance protein found in *Enterobacteriaceae*) (CARD) (39). The presence of all these resistance genes in *E. coli* from cattle manure is not unexpected since tetracyclines, beta-lactams, cephalosporins, aminoglycosides, trimethoprim, or quinolones are used to treat various infections in cattle, including respiratory diseases, mastitis or dry cow therapy (42).

Recently, a huge healthcare problem arose following infections caused by ESBL-producing bacteria, with the CTX-M producers predominating. Among them, the most widespread seems to be CTX-M-15, belonging to the CTX-M-1 group, followed by CTX-M-14 (43, 44). Most of the known ESBL enzymes are expressed by genes mapped on plasmids (45). Moreover, *E. coli* has been recently recognized as a major source of ESBLs during nosocomial and community-associated infections (44, 46).

Plasmid-encoded genes can arise from multiple sources and be further disseminated by HGT. Our present findings indicate the presence of beta-lactam-resistance genes originating in dairy cattle in various MGEs, such as insertional sequences or transposons (Fig. 4A-D). ESBL-producing *E. coli* is commonly isolated from community or hospital infections, human fecal carriage, and to an increasing degree from food-producing, companion and wildlife animals, and the environment; however, particular plasmids similar to those found during this study, have been found only in clinical samples (47).

Our findings confirm previous observations that ESBL transmission is mainly driven by insertion sequences, transposons, integrons, and plasmids, which are homologous and may be isolated from both food-producing animals and humans (45). The wide spread of *E. coli* carrying the *blaCTX-M-15* gene seems to be linked to a hot spot of insertion on the *blaTEM-1*-Tn*2* transposon, which is largely found in broad-range host spectrum IncF plasmids (47). It has also been noted that plasmids expressing an ESBL phenotype also frequently carry genes encoding resistance to other commonly-used antimicrobial drug classes, such as aminoglycosides, chloramphenicols, fluoroquinolones, or tetracyclines (45, 48); this was also noted during this study. Plasmid-mediated transfer of drug-resistance genes between various bacterial species is considered to be one of the most important mechanisms driving the spread of multidrug resistance (45).

By definition, plasmids do not carry genes essential for the growth of host cells under non-stressed conditions: plasmids frequently contain a cargo of genes involved in adaptation to environmental challenges, such as ARGs (49–52). Plasmids represent an important pool of adaptive and transferable genetic elements, with large plasmids (>30 kb) being commonly capable of transferring their genetic information between bacteria (53). HGT facilitates rapid phenotypic evolution, the incorporation of new metabolic pathways and network expansions, and can eventually lead to ecological speciation across microbes; however, to be maintained at high frequencies within the population, the regions transferred by HGT must provide fitness benefits and minimize fitness cost (54). The magnitude of fitness costs for any evolutionary event, including HGT, depends strongly on how the environment structures its selection pressures. It is clear that selection efficiently entails even the most subtle of costs (55). Removing a factor responsible for selection pressure will result in the loss of a plasmid carrying genes that determine survival in unfavorable conditions; however, the plasmids can remain within the cell when it harbors more than one resistance determinant. Thus, even though some antibiotics were not applied during therapy, the bacteria can still carry identified plasmids due to cross-resistance (56).

Although the particular plasmids identified during the study have not been described previously, similar plasmids (threshold 0.99) have been reported both in countries with restrictive antimicrobial usage policies (Denmark), and in those with ’flexible’ ones (India, China). Resistance plasmids have been found across the globe (USA, Europe, Asia), which implies that they are also widespread among people and the environment; indeed, they have been isolated from human clinical samples (urine, blood) and river sediments. No similar plasmids have previously been isolated in Poland, and none have originated from dairy cattle. The accumulation of so many ARGs on a single plasmid may result from the huge selection pressure placed on bacteria living in the intestines of dairy cattle due to the overuse of antibiotics in dairy cattle breeding (12).

The newly-identified plasmids were included in the IncY incompatibility group, which unlike IncN or IncP, is not a broad host-range incompatibility group. The prevalence of IncY plasmids seems to be limited to the *Enterobacteriaceae* family. Similar plasmids (threshold 0.99) have previously been identified in *E. coli*, *Klebsiella pneumoniae,* and *Salmonella enterica subsp. enterica*; however, other incY plasmids have also been reported in *Cronobacter sakazakii*, *Escherichia fergusonii*, *E. albertii, E. marmoae, Enterobacter hormaechei, E. cloace, Shigella flexneri,* and *Klebsiella aerogenes*. Interestingly, these other incY plasmids were isolated from a wide range of sources, ranging from human clinical samples, wild animals (crow, wild boar), companion animals (dog), livestock (pig, cattle, sheep), poultry, and environmental samples (rivers or wastewater treatment plants), to even alfalfa sprouts (36, 37). In addition, incY plasmids have been found in meat and other food samples. Balbuena-Alonso et al. (2) report that plasmids originating in food contaminated with *E. coli* may be transferred to pathogenic strains of *E. coli*.

Our findings have considerable clinical and environmental implications. The presence of MDR bacteria in various settings poses significant challenges to public health. Research emphasizes the need for robust antibiotic stewardship and environmental monitoring to manage the spread of resistance. The cost of treating a single infection caused by MDR bacteria is 165% higher than for non-MDR infection, with an incremental cost of 1383 USD (57). Highly resistant Gram-negative infections were estimated to require between 39 and 138.2 days of therapy for 10,000 patient encounters. In other countries, rates are even higher, with over 10% of Gram-negative bacteremia caused by difficult-to-treat resistant pathogens (58).

Antibiotic-resistant infections are a major public health concern around the world. Recent data show substantial increases in the use of vancomycin and broad-spectrum antibiotics such as carbapenems, as well as third- and fourth-generation cephalosporins, and β-lactam/β-lactamase inhibitor combination antibiotics. These data, combined with evidence that 30% of antibiotic prescriptions may be inappropriate, suggest that antibiotic-resistant bacteria will soon represent a substantial threat (59).

It is crucial that antibiotic-resistant bacterial strains and the plasmids coding for resistance are tracked effectively. Antibiotic resistance does not respect boundaries, and ARGs can move between bacterial strains in an almost uncontrolled manner. *E. coli* demonstrates considerable diversity and high adaptability, and hence can inhabit the natural environment, animal intestines, and plants, where it can serve as both a harmless commensal and a dangerous pathogen. As such, this bacterium an ideal vector for transmitting plasmids encoding drug resistance between different environments.

## Conclusion

The study characterizes two *E. coli* plasmids, each carrying ARGs and insertion elements, found in two representative isolates from dairy cattle manure. Such plasmids have not been previously reported in Poland, though similar ones have been recorded in crucial human pathogens in other countries. However, plasmids resembling pECmdr1.1 and pECmdr2.1 from cattle manure-derived *E. coli* have only been seen in aquatic environments and human clinical samples, not in farm animals.

The identified plasmids, isolated from *E. coli* obtained from cattle manure, confer resistance to β-lactam antibiotics, including penicillins (even with β-lactamase inhibitors like clavulanic acid), cephalosporins (second to fourth generation), aminoglycosides, fluoroquinolones, tetracycline derivatives (glycylcycline antibiotics), and chemotherapeutics / sulfonamides (trimethoprim / sulfamethoxazole). All identified resistance plasmids carry conjugal transfer genes, enabling HGT. The identified plasmids belong to the IncY group, which have been identified in *Enterobacteriaceae* such as *E. coli*, *Salmonella* spp., and *Klebsiella pneumoniae*, though information on IncY plasmids remains limited.

## Author contributions

M.Z. - Methodology, Formal analysis, Investigation, Data Curation, Writing - Original Draft, Visualization; A.B. - Formal analysis, Investigation; J.G. - Formal analysis, Data Curation; A.D., S.M., K.G - Sequencing and primary Data Curation; P.K – Investigation, Data Curation; M.P. - Conceptualization, Writing - Review & Editing, Project administration, Funding acquisition.

## Data availability

The genome sequences of *E. coli* strains generated during this study have been deposited in GenBank (NCBI) with the accession number PRJNA942482. Plasmids datasets generated during this study have been deposited in the University of Warsaw repository with the doi number doi:10.58132/YFTW53.

